# LEAF SHAPE OF QUINOA’S WILD ANCESTOR *CHENOPODIUM HIRCINUM* IN A GEOGRAPHIC CONTEXT

**DOI:** 10.1101/2024.08.21.608958

**Authors:** Ramiro N. Curti, Jonatan Rodriguez, Pablo Ortega-Baes, Sergio J. Bramardi, Eric Jellen, David E. Jarvis, Peter J. Maughan, Mark Tester, Héctor D. Bertero

## Abstract

**Premise:** Intraspecific variation in plant traits, such as leaf morphology, offers insights into local adaptation and the ecological niche breadth of species. *Chenopodium hircinum*, the wild ancestor of quinoa, is widely distributed across various ecoregions in Argentina. A detailed comparative analysis of leaf morphology across 23 populations covering its entire distribution was conducted to understand patterns of intraspecific variation along geographic gradients.

**Materials and Methods:** Using 104 leaves from these populations, leaf shapes were described through shape, landmarks, and Fourier descriptors. Both univariate and multivariate analyses were employed to evaluate variations among and within populations and to link leaf shape patterns with climatic and geographical variables at the sites of origin.

**Results:** A significant variability in leaf shapes was found across all ecoregions, with marked integration among populations. While landmark-based and Fourier analyses differentiated populations, shape descriptors introduced complexity in identification. Leaf morphology varied widely, from rounded and toothed to sharply lobed, with prominent lobes located at the base or middle of the leaf. The key environmental factors influencing leaf shape, particularly lobulation and relative positioning, were mean annual temperature and altitude.

**Conclusions:** The findings suggest that in *C. hircinum*, leaf shape variability goes beyond the simple lobed vs. entire dichotomy and is shaped by environmental temperatures. Leaf morphology and margin traits are primarily governed by the relative position of the lobes, which are strongly correlated with the mean temperature at the population’s origin. This convergence across different ecoregions highlights the need for further research to explore the functional significance of such variation.

---

Leaf shape is a crucial trait in plants, as it influences processes such as photosynthesis, transpiration, and thermal regulation (Givnish and Kriebel, 2017). Shape quantifies leaf form in terms of the natural dimensions or size (Nicotra et al., 2011). Several studies have demonstrated a convergence between climate and leaf morphology, with smaller, lobed leaves being predominant in colder climates and larger, entire leaves more common in warmer climates (Royer et al., 2005, 2012; Traiser et al., 2005; Peppe et al., 2011; Yang et al., 2015; Li et al., 2020). This variation reflects the need to optimize water and thermal efficiency in different environments (Royer and Wilf, 2006). Differences in leaf shape between temperate and tropical climates, for example, are explained by adaptation to water availability and temperature (Fritz et al., 2018; Gleason et al., 2018). Sensitivity to these factors varies among tree species, shrubs, lianas, and herbaceous plants, with a more pronounced response observed in woody plants due to their structure and longevity (Traiser et al., 2005; Frye et al., 2020).

The insensitivity of leaf margin traits in herbaceous species to regional climate is attributed to their adaptation to subcanopy microclimates (Royer et al., 2012). However, Hightower et al. (2024) recently showed a strong relationship between leaf shape variation and temperature in weedy specimens’ from *Capsella bursa-pastoris* (Brassicaceae) collected throughout the continental U.S. over a long period. Accordingly, the lack of sensitivity in herbaceous plants suggests that it is not a universal phenomenon, highlighting the need for alternative model systems to study within-species leaf shape variation across diverse environmental conditions. One promising model system is the use of wild relatives of crops, as these herbaceous species hold critical genetic resources for improving crop resilience under future climate change scenarios (Renzi et al., 2022). These wild relatives are typically found in marginal habitats and have evolved in situ, adapting to the increasingly adverse effects of climate change (Maxted et al., 2012). Elucidating the variation in traits that allow these plants to thrive in extreme environments presents a critical challenge for researchers globally. This understanding is essential for enhancing crop gene pools with advantageous traits derived from wild relatives (Bohra et al., 2022).

*Chenopodium hircinum* (Amaranthaceae), also known as avian goosefoot, is the likely wild ancestor of quinoa (*C. quinoa*) (Jarvis et al., 2017; Maughan et al., 2019), a pseudocereal crop that recently gained international attention because of its nutritious seeds (Bertero, 2021). This species is widely distributed throughout temperate, subtropical, and highland South America, with a high proportion of its geographic range in Argentina (Wilson, 1990; Curti et al., 2023). In Argentina, avian goosefoot can be found in the northwest Andean uplands, eastward into the humid Pampas, and southward into the Patagonian plains (Wilson, 1988). Within this extensive geographic range, avian goosefoot populations cover several ecoregions characterized by contrasting environmental conditions, with seasonal and diurnal temperature variation, relative humidity, and precipitation gradients increasing from the west to the east (Curti et al., 2023).

Current research on avian goosefoot focuses on uncovering genetic variation related to heat stress tolerance, as recent studies indicate that quinoa performs poorly in environments with temperatures exceeding 35°C (Lesjak and Calderini, 2017; Hinojosa et al., 2018). A previous study on avian goosefoot populations in Argentina revealed significant genetic variation in phenological traits across accessions collected from different geographic regions, largely associated with temperature gradients throughout its distribution (Curti et al., 2022). Given that leaves are the primary sites of photosynthesis and thermoregulation, it is expected that variations in phenological and leaf traits will covary across populations from different climatic zones and ecoregions.

In this species, leaves are defined by trilobed marginal dissection; however, large variability is found in plants from different geographic sites (Wilson, 1988). An early study describing leaf blades did not differentiate populations of avian goosefoot distributed within Argentina (Wilson, 1988). In turn, these avian goosefoot samples were differentiated from Andean samples allied to *ashpa*-*quinua-*complex (Wilson, 1988). However, populations sampled by Wilson (1988) did not cover the entire geographic range in Argentina.

In this context, we have studied the leaf shape of Argentinean avian goosefoot using several morphometric tools. The populations sampled cover most of the species-wide geographic range in the country, with clear temperature and precipitation gradients increasing from the west to the east. We believe that analysis of avian goosefoot populations from contrasting origins within its geographic range should enable us to determine whether leaf shape is predictive of climatic conditions and ecoregion types. If leaf shape is associated with climatic conditions or ecoregions, then population differences should associate with specific ecoregions. In the present study we answer the following questions: (1) How phenotypically different are avian goosefoot leaves at ecoregional and intraspecific levels? (2) Does the leaf shape of avian goosefoot populations vary along the geographic range of Argentina? (3) Is geographic variation in leaf shape related to ecoregions and their climatic variables at the provenance sites?

## MATERIALS AND METHODS

### Collection sites and climatic data

In this study, seeds from 23 populations of avian goosefoot collected in Argentina during February and March of 2017 were analyzed. These populations were sampled from a range of geographical coordinates: latitudes from 24°89′S to 35°25′S, longitudes from 59°21′W to 68°33′W, and altitudes ranging from 32 to 2,116 meters above sea level, across eight provinces of Argentina (Appendix S1, Supplemental Data). The populations were categorized according to the ecoregions defined by Olson et al. (2001): High Monte (HM), Southern Andean Yungas (SAY), Dry Chaco (DC), Low Monte (LM), and Humid Pampas (HP) (Appendix S1).

Geographical coordinates from each collection site were used to extract climatic data, including Mean Annual Temperature (MAT, °C) and Annual Precipitation (AP, mm), from the WorldClim website (https://www.worldclim.org/data/index.html). These data were obtained at a spatial resolution of 30 arc-seconds latitude and longitude (Appendix S1).

### Common garden experiment and leaf sampling

A common garden experiment was conducted in a greenhouse at the School of Agronomy, University of Buenos Aires, to assess variability in phenological traits among avian goosefoot populations. Detailed descriptions of the experimental growing conditions can be found in Curti et al. (2022). In total, 104 fully-expanded leaves were sampled from three to five individuals per population (Appendix S1). Leaves were collected at the anthesis stage, when the flowers of the main panicle open and anthers extrude pollen (Curti et al., 2022). Specifically, leaves were harvested from the middle third of the main stem of each plant. This sampling method was chosen because leaf marginal dissections are fully developed at anthesis (Ruiz and Bertero, 2008) and this position is commonly used for morphological descriptions of quinoa varieties and related wild species (International et al., 2013). After the petioles were removed, leaves were scanned using an Epson L355 scanner at a resolution of 300 dots per inch (dpi) and saved as standard JPEG files for subsequent image analysis.

### Shape descriptors, landmarks and outline analysis

Leaf images of each individual were converted into black silhouettes, and several shape descriptors (hereafter referred to as “shp”) were calculated using ImageJ (https://imagej.net/ij/). These descriptors included: circularity calculated as 4π^2^(area/perimeter^2^)), which measures the ratio of the leaf area to the perimeter outline; aspect ratio, the ratio of the major axis to the minor axis of the leaf; solidity the ratio of the leaf area to the area of the convex hull surrounding the leaf; basal angle (Bangle) and top angle (Tangle). These angles describe the leaf’s shape at its basal and apex regions, respectively.

Eight landmarks (hereafter referred to as “ldks”) were defined to outline the leaf shape of *C. hircinum* using the ImageJ point tool. The landmarks were positioned as follows: first and second ldks at the tips of the lobes with the largest width of the leaf blade; third and seventh ldks at the tips of the main lobes on each side of the leaf blade; fourth and sixth ldks at the sinus valleys below the main lobes; fifth and eighth ldks at the petiolar junction and the apex of the leaf blade, respectively.

Elliptical Fourier Descriptors (EFDs) were calculated to describe the leaf outlines. This was done by progressively decomposing the outline coordinates (*x*, y) into harmonics, as detailed by Bonhomme et al. (2014). Prior to analysis, the leaf outlines were normalized with the petiolar junction as the starting point for each outline.

### Statistical analysis

Multivariate analyses of variance (MANOVA) were employed to assess significant differences between Ecoregions and Populations nested within Ecoregions for shp, ldks, and EFDs using the PROC GLM in SAS© software (SAS Institute, 2004). In addition, separate analyses of variance (ANOVA) were conducted for each trait. For both the MANOVAs and ANOVAs, a linear mixed-effect model was applied, with Ecoregion treated as a fixed effect and the nested factor considered as a random effect. Due to the imbalance in the dataset, hypothesis tests for all effects were based on appropriate error terms derived from the expected mean squares. For shp, the MANOVA was conducted using transformed values of circularity (1/circularity), aspect ratio (1/aspect ratio), and solidity (1/solidity) to satisfy the assumption of multivariate normality. A scatterplot matrix was created to examine associations between pairs of shape descriptors. For ldks and EFDs, the MANOVAs were based on principal components (PCs) obtained from previously conducted Principal Component Analyses. The PCs for both methods were derived following a generalized Procrustes analysis using the ‘shapes’ (Dryden and Mardia, 2016) and ‘Momocs’ (Bonhomme et al., 2014) packages, respectively. The selection of the relevant number of harmonics to describe leaf shape was guided by the harmonic power test, while the determination of the appropriate number of PCs was informed by broken-stick models using the ‘PCDimension’ package (Wang et al., 2018).

To investigate the influence of climate and geographical variables on leaf shape variation, we performed a Redundancy Analysis (RDA) using the ‘vegan’ package (Oksanen et al., 2001). In this analysis, the shape descriptors and PCs of ldks and EFDs that explained significant variation in leaf shapes were treated as response variables. The predictor variables included latitude, longitude, and altitude (geographical variables), as well as MAT and AP (climate data). We conducted a permutation test with 1000 iterations to assess the significance of the overall model, individual predictor variables, and the axes. A biplot was generated to simultaneously visualize the relationships among the response and explanatory variables.

## RESULTS

### Shape descriptors, landmarks and outlines analyses

Based on Pearson correlation tests, the analysis revealed that leaf circularity showed a positive correlation with bangle, tangle, and solidity. Conversely, leaf circularity had a negative correlation with aspect ratio (Appendix S2). Additionally, the aspect ratio was found to be negatively correlated with bangle, meanwhile, solidity exhibited a positive correlation with tangle (Appendix S2).

The broken-stick model indicated that only the first two PCs were significant for capturing the variance in the data, together accounting for 62.5% of the total variance (Figure 1). PC1 explained 38.8% of the variance and was associated with variations in the distances between specific ldks on the leaf. This component described distances between basal ldks 1 and 8, and 2 and 8; apex ldks 4 and 5, and 5 and 6; and between ldks 3 and 4, and 6 and 7 (corresponding to sinus valleys and tips of the main lobes) (Figure 1). High values on PC1 were associated with leaves that have rounded shapes, characterized by a short distance between basal and apex ldks, and deep basal lobing, as indicated by increased distances between sinus valleys and tips of the main lobes (Figure 1). PC2 accounted for 23.8% of the variance and reflected differences in distances between ldks 1 and 7, and 2 and 3; as well as between ldks 3 and 4, and 6 and 7 (sinus valleys and tips of the main lobes) (Figure 1). Higher values on PC2 corresponded to leaves with a larger proportion of the blade concentrated in the middle, characterized by greater distances between ldks 1 and 7 and 2 and 3, and pronounced mid-lobe depth (Figure 1). This configuration helps differentiate populations based on their leaf morphology, particularly in relation to mid-lobe prominence and overall shape.

**Figure 1.**
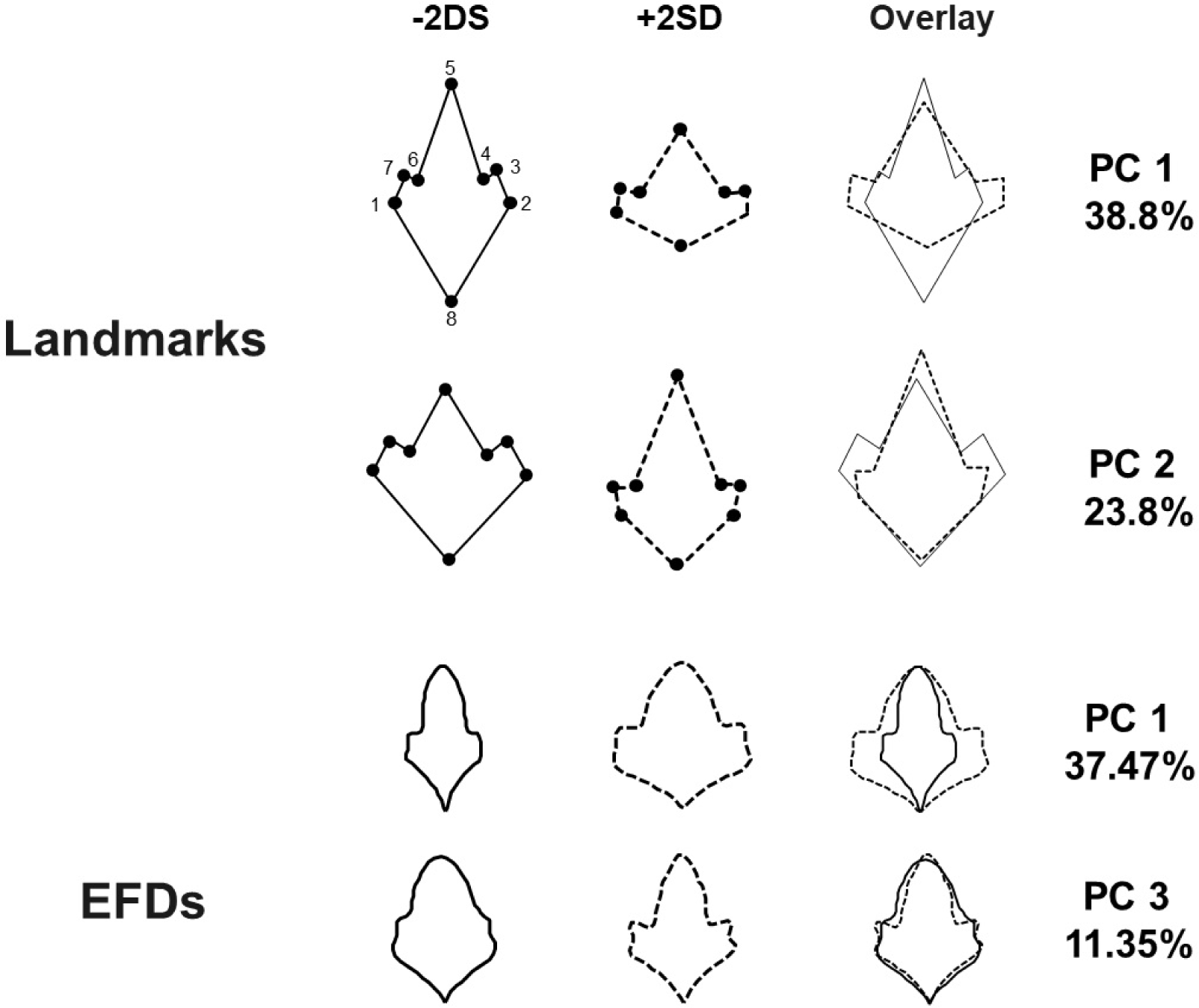
Eigenleaves generated from a PCAs applied to landmarks and Elliptical Fourier Descriptors (EFDs). The significant principal components (PCs) contributing to shape description are presented alongside the percentage of variance they explain. For each PC, the eigenleaves at −2 standard deviations (SD) (solid line) and +2 SD (dashed line) along the PC axis are illustrated. The overlay of eigenleaves at ±2 SD highlights the shape variance captured by each PC. The positions of eight key landmarks are also displayed.

A total of 17 harmonics were selected to reconstruct the leaf shapes (Appendix S3). The broken-stick model identified the first five harmonics as significant, together accounting for 79.81% of the total variance. PC1 explained 37.47% of the variance and was associated with changes in basal lobing and overall leaf area. Specifically, PC1 differentiated between leaves with deep basal lobing and a large basal area on the right side and leaves with shallow basal lobing and a more constrained basal area on the left side (Figure 1). PC3 and PC5 accounted for 11.35% and 4.82% of the shape variance, respectively. PC3 was linked to variations in the size of the middle leaf areas, while PC5 was associated with the development of the main lobes of the leaf (Figure 1). These components highlight specific aspects of leaf morphology related to the middle region and the prominence of the main lobes.

### Variation among and within populations

The MANOVA results indicated that the multivariate means across Ecoregions were not significantly different for shp (F_20, 47.83_ = 1.21; P = 0.28), ldks (F_12, 42.62_ = 0.86; P = 0.594), and EFDs (F_12, 47.83_ = 1.11; P = 0.374). However, the term for Populations nested within Ecoregions was statistically significant for all morphometric methods: shp (F_90, 378.01_ = 4.49; P less than 0.001), ldks (F_54, 236.21_ = 5.51; P less than 0.001), and EFDs (F_90, 378.06_ = 6.16; P less than 0.001). Separate ANOVAs confirmed that variation among Ecoregions was not significant for any of the morphometric methods (Table 1). Instead, a substantial portion of the variation was attributed to both nested populations and residual terms (Table 1). For most shape traits, except for tangle, the variance components were nearly equally distributed between and within populations (Table 1). Conversely, for ldks and EFDs, the variation among populations generally exceeded the variation within populations for the retained PCs (Table 1).

**Table 1:** Results of separate ANOVAs for each trait using respective morphometric methods. Mean squares are shown for the fixed effect (Ecoregion), while the variance component ± standard errors for nested and residual sources of variation. The percentages of variance accounted for random effects are shown in parentheses next to each variance component.

| Morphometric method | Traits | Ecoregion | $\sigma^2_{Population(Ecoregion)}$ | $\sigma^2_{Residual}$ |
| --- | --- | --- | --- | --- |
| Shape descriptors | 1/Circularity | 0.61ns | 0.07±0.03 (40.72)** | 0.09±0.01 (48.82)** |
| | 1/Aspect ratio | 0.01ns | 0.01±2.40e-3 (48.35)** | 0.01±1.06e-3 (51.65)** |
| | 1/Solidity | 0.01ns | 1.61e-3±6.60e-4 (42.90)** | 2.00e-3±3.14e-4 (53.38)** |
| | Bangle | 130.26ns | 60.70±25.66 (38.85)** | 95.55±15.10 (61.15)** |
| | Tangle | 2808.47ns | 608.89±198.20 (74.38)** | 209.65±32.97 (25.61)** |
| Landamarks | PC1 | 2.45ns | 0.71±0.23 (70.82)** | 0.29±0.04 (29.17)** |
| | PC2 | 3.91ns | 0.52±0.20 (48.55)** | 0.55±0.01 (51.44)** |
| Elliptical Fourier Descriptors | PC1 | 2.45ns | 1.59e-3±5.22e-4 (72.08)** | 6.17e-4±9.70e-5 (27.92)** |
| | PC3 | 4.09e-3ns | 3.45e-4±1.36e-4 (59.59)** | 2.34e-4±3.70e-5 (40.41)** |
| | PC5 | 9.24e-4ns | 1.87e-4±6.70e-5 (65.85)** | 9.70e-5±1.50e-5 (34.15)** |

\* Significant at $P < .05$ ; \*\* at $P < .01$ ; ns: nonsignificant.

### Relationship between leaf shape and environmental variables

The results RDA indicated that environmental variables explained 20% of the variation in leaf shape across populations. The model was statistically significant (F_5, 908_ = 4.91; P less than 0.001), with all predictor variables contributing significantly to the explained variation. Among the RDA components, only the first three explained a significant portion of the variation (Appendix S4). In the biplot, leaf shape variation was predominantly associated with MAT on the first RDA component, and with geographical variables on the second component (Figure 2). The AP had a minor impact on the variation, as indicated by the short length of its vector. The individuals from all populations were dispersed throughout the Euclidean space, showing a wide range of leaf shapes (Figure 2). Specifically, populations with leaf blades featuring large basal areas and deep lobes were located on the right side of RDA1, corresponding to colder, high-altitude sites (Figure 2). In contrast, populations with elongated leaf blades and prominent lobes in the middle regions were positioned on the left side of RDA1, associated with warmer sites at higher latitudes or longitudes along RDA2 (Figure 2).

**Figure 2.**
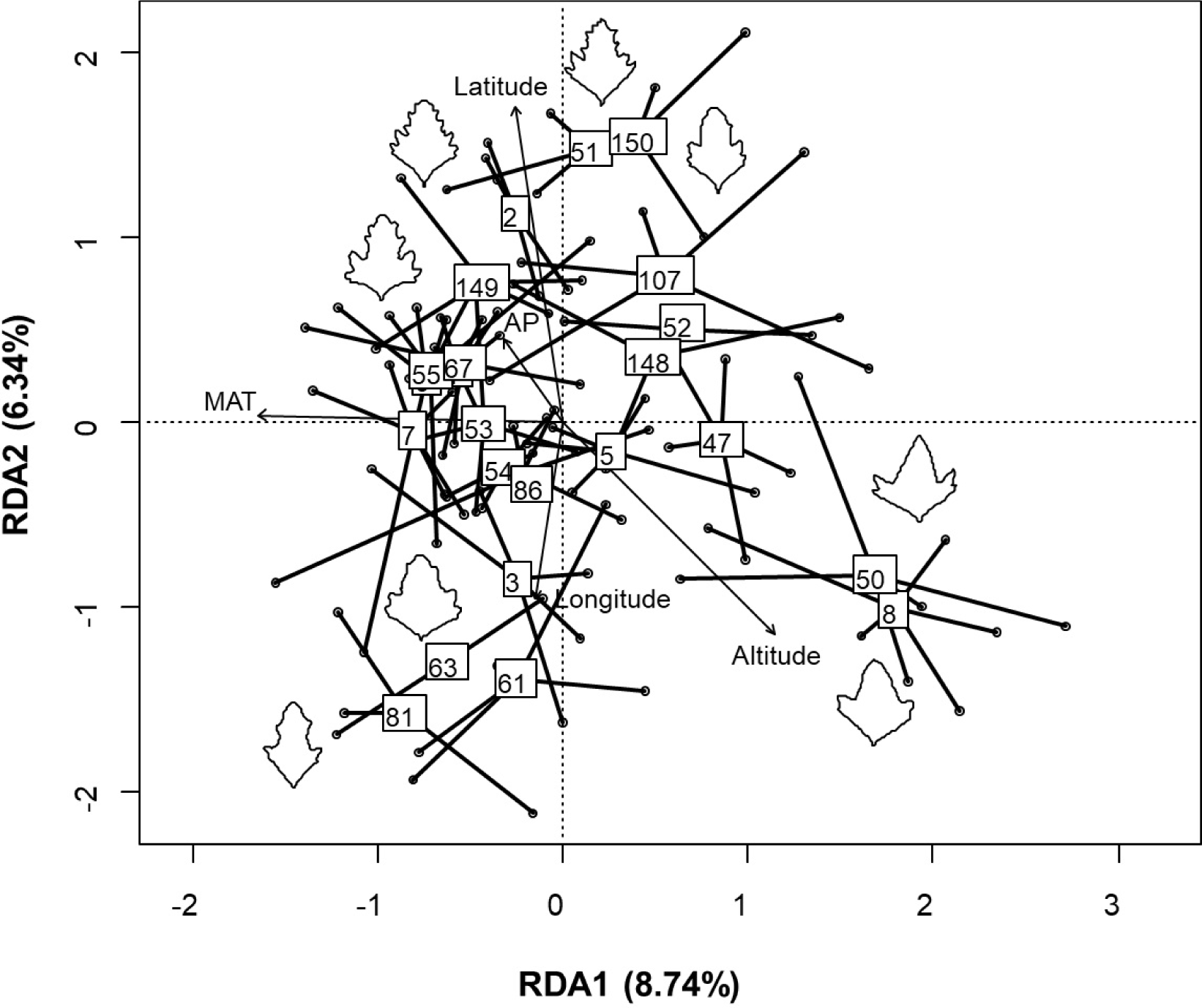
Biplot with factor scores derived from redundancy analysis, with explanatory variables represented as vectors (arrows), MAT: mean annual temperature and AP: annual precipitation. Populations are indicated by boxes, with individual coordinates connected to their respective population centroids. Representative leaves from populations occupying the extreme positions along the axes are also displayed.

## DISCUSSION

We found that populations of avian goosefoot from Argentina exhibit significant variations in leaf shape across their geographical distribution. This extensive variation was influenced more by differences at the inter- and intrapopulation levels than by ecoregion differences (Table 1). However, the extent of variation attributable to among and within population levels varied depending on the morphometric methods used (Table 1). The analysis revealed that substantial variation within populations in almost all shape descriptors indicates that each population contains a diverse range of leaf types. In contrast, both landmark-based and Fourier analysis methods were effective in capturing a significant proportion of the variation among populations (Table 1). These methods highlighted clear distinctions in leaf shape between populations, despite the overall high variability within individual populations.

Leaf shape among our sampled populations of avian goosefoot exhibited considerable variation, ranging from rounded and toothed to sharply pointed, with lobed blades at the basal and middle thirds (Figure 2). This variation was primarily observed in the apex rather than the basal angle. A similar combination of leaf shape traits distinguishes the diploid species *C. watsonii*, *C. fremontii*, *C*. *neomexicanum*, and *C*. *palmeri*, which inhabit the western North America (Walters, 1988). However, despite their wide geographic distribution, leaf shape variation among these species did not exhibit a clear ecotypic structure. Indeed, psammophytic New World A-genome diploids, which have been shown to be closely related to the *C. berlandieri*-*hircinum*-*quinoa* complex (Young et al., 2023), are well known to have narrow leaves, a presumed adaptation to water scarcity. These include North and South American *C. desiccatum* and *C. pratericola*; North American *C. cycloides*, *C. leptophyllum*, *C. littoreum*, *C. pallescens*, and *C. subglabrum*; and South American *C. papulosum*. Variation in lobed blades at the basal and middle thirds was recently shown to differentiate subspecies of *C*. *oahuense* (Bullock and Cantley, 2022), an allopolyploid derivative of central and east Asian *C*. *acuminatum* and the ancestor of *C*. *suecicum* (Mandák et al., 2016). *C*. *suecicum* played a role in the early hybridization stage of the allotetraploid complex that eventually gave rise to domesticated *Chenopodium* species in the New World (Jarvis et al., 2017). These findings suggest that leaf shape variation among *Cellulata* species associated with quinoa and its wild relatives has a deep evolutionary history, tracing back to their radiation across Eurasia and the Americas.

Our results indicate that populations from all ecoregions were intermixed, exhibiting minimal differentiation based on ecoregional structure. This finding is consistent with Wilson’s (1988) earlier study, which identified a coherent grouping of eastward and upland Argentinean populations of avian goosefoot. The observed variation pattern was anticipated, as the ecoregions encompass a broad geographical range with diverse environmental conditions that create transition zones (Olson et al., 2001; Oyarzabal et al., 2018). Although there is a lack of research specifically addressing leaf shape variation among and within populations of herbaceous plants in Argentina, a comparable range of intraspecific variation has been documented in wild relatives of sunflower (*Helianthus annuus*) that have naturalized in various geographic regions of Argentina (Presotto et al., 2009).

Mean annual temperature and altitude were the primary geographic and climatic factors influencing the ordination of avian goosefoot populations along the first axis of the Redundancy Analysis (RDA), while latitude and longitude contributed to the second axis (Figure 2). These findings partially align with previous research indicating that latitude is the key geographic factor distinguishing southern and northern populations of avian goosefoot (Wilson, 1988). However, our results further suggest that leaf shape variation is primarily associated with temperature gradients within the species’ geographic range. This geographic pattern of variation was previously observed in phenological traits (Curti et al., 2022). Since leaf shape and phenological traits tend to covary similarly across the environmental gradient, it is reasonable to assume that a high degree of phenotypic integration is maintained (Westerband et al., 2021). This suggests not only that selection should mold the responses of individual traits to temperature variation in Argentinean avian goosefoot populations, but that it should also act on the interrelationships among traits at plant level.

The extent to which the broad phenotypic plasticity observed among and within populations of *C*. *hircinum* results from genetic variation, environmental influences, or a combination of both remains a subject of investigation. Wilson’s (1988) early study revealed substantial variation in isozyme markers, which was not consistent with the conservative phenotype observed in leaf and seed traits. While Wilson (1981) attributed this pattern to similar selective pressures in a relatively uniform habitat, our results suggest that this species’ wide phenotypic plasticity enables it to adapt to heterogeneous environmental gradients, particularly temperature variations across its broad geographic range. Future studies should evaluate to what extent the structure of phenotypic correlations among leaf shapes and phenological characters resulted from genetic correlations developed under specific temperature conditions.

The pattern of variation in leaf shape observed across temperature gradients in this study differ with general trends documented in several tree, vines and shrubs species (Peppe et al., 2011; Royer et al., 2012). Specifically, leaf shape in Argentinean avian goosefoot populations varied from rounded-toothed to sharpened with strong basal or middle lobes, rather than displaying a simple distinction between entire vs. lobed forms. Variation in basal and middle lobing was particularly associated with sites differing in MAT, suggesting that this trait responds to temperature gradients. Populations inhabiting high-elevation sites with lower MAT exhibited leaves with pronounced marginal dissections and strong basal lobes (Figure 2). In contrast, populations from plains with higher MAT displayed leaves with lobes concentrated in the middle area (Figure 2). This suggests that the variation in the relative position of lobing among Argentinean avian goosefoot populations is likely associated with to temperature differences. Future research is needed to unravel the functional significance of differential lobulation along the leaf blade in populations from contrasting environments.

## CONCLUSIONS

Leaf shape in avian goosefoot exhibits a broad range of morphological diversity across its native range in Argentina. The variation spans from rounded-toothed to sharp-edged leaves with pronounced basal or middle lobes, predominantly observed within populations rather than across ecoregions. This variation is significantly correlated with the mean growing season temperature, indicating a substantial influence of regional climate on leaf morphology. These findings enhance our understanding of phenotypic diversity in quinoa’s wild ancestor and offer insights into how intraspecific variation in avian goosefoot may influence its ecological niche breadth.

## AUTHOR CONTRIBUTIONS

R.N.C., E.J., D.E.J. and H.D.B. collected seeds of *C*. *hircinum* populations from Argentina. R.N.C., P.O.B., and H.D.B. designed the study. H.D.B. grew plants in the greenhouse and conducted the common garden experiment. R.N.C., J.R., S.J.B., and H.D.B. conducted all data analyses. H.D.B. and M. T. Resources and Funding acquisition. All authors contributed to writing and editing the manuscript and approved the final submission.

## ACKNOWLEDGMENTS

This research was supported by FONCYT (Fondo Nacional de Ciencia y Técnica, project PICT-2018-03456) and CONICET (the Argentine Scientific Research Council, project PIP 11220170100459 CO). The *C*. *hircinum* collection trip was supported by grant OSR-2016-CRG5-2966 from KAUST University.

## DATA AVAILABILITY STATEMENT

The data that support this study will be shared upon reasonable request to the corresponding author.

---

**APPENDIX S1.** Passport data of Argentinean populations of *C*. *hircinum* with geographic and climatic data of collection sites described by morphometric analyses.

| Code | Pop | Site | Province | Ecoregion | Samples | Lat (S) | Lon (W) | Altitude | AP | MAT |
| --- | --- | --- | --- | --- | --- | --- | --- | --- | --- | --- |
|  |  |  |  |  | *n* |  |  | (masl) | (mm) | (°C) |
| 1 | P01 | Mabran | Santiago del Estero | DC | 5 | 29.35 | 62.44 | 83 | 816 | 19.93 |
| 2 | P02 | Real Sayana | Santiago del Estero | DC | 5 | 28.82 | 62.85 | 98 | 701 | 20.63 |
| 3 | P03 | Cerrillos | Salta | DC | 5 | 24.89 | 65.47 | 1231 | 688 | 16.88 |
| 4 | P05 | Rosario de la Frontera | Salta | DC | 5 | 25.81 | 64.98 | 790 | 641 | 18.88 |
| 5 | P06 | Trancas | Tucumán | DC | 4 | 26.24 | 65.28 | 775 | 452 | 19.44 |
| 6 | P07 | Benjamín Paz | Tucumán | DC | 5 | 26.37 | 65.30 | 769 | 488 | 19.49 |
| 7 | P08 | Tafí del Valle | Tucumán | SAY | 5 | 26.90 | 65.69 | 1913 | 460 | 14.88 |
| 8 | P47 | Tolombón | Salta | HM | 5 | 26.23 | 65.94 | 1660 | 211 | 16.03 |
| 9 | P50 | San José | Catamarca | HM | 5 | 26.78 | 66.06 | 1956 | 213 | 15.30 |
| 10 | P51 | Desmonte | Catamarca | HM | 4 | 26.88 | 66.10 | 2063 | 211 | 15.04 |
| 11 | P52 | Punta de Balasto | Catamarca | HM | 3 | 26.96 | 66.14 | 2116 | 213 | 14.96 |
| 12 | P53 | Los Nacimientos | Catamarca | HM | 5 | 27.16 | 66.72 | 2064 | 149 | 15.58 |
| 13 | P54 | Hualfín | Catamarca | HM | 5 | 27.22 | 66.84 | 1869 | 146 | 16.25 |
| 14 | P55 | Belén | Catamarca | HM | 5 | 27.63 | 67.03 | 1355 | 172 | 17.69 |
| 15 | P61 | Condorhuasi | Catamarca | HM | 5 | 27.58 | 65.99 | 1809 | 378 | 16.73 |
| 16 | P63 | Singuil | Catamarca | SAY | 3 | 27.82 | 65.86 | 1206 | 503 | 17.91 |
| 17 | P67 | Rio Los Puestos | Catamarca | SAY | 5 | 28.04 | 65.82 | 1054 | 492 | 18.28 |
| 18 | P81 | Sumampa | Santiago del Estero | DC | 3 | 29.39 | 63.47 | 241 | 742 | 19.38 |
| 19 | P86 | Sumampa | Santiago del Estero | DC | 5 | 29.39 | 63.47 | 241 | 742 | 19.38 |
| 20 | P107 | Dean Funes | Córdoba | DC | 5 | 30.41 | 64.38 | 661 | 616 | 16.88 |
| 21 | P149 | Ceres | Santa Fe | DC | 3 | 34.58 | 68.56 | 864 | 332 | 14.13 |
| 22 | P148 | 25 de Mayo | Mendoza | LM | 5 | 29.86 | 61.95 | 89 | 895 | 19.18 |
| 23 | P150 | Roque Pérez | Buenos Aires | HP | 4 | 35.42 | 59.36 | 32 | 988 | 15.99 |

Abbreviations: DC: Dry Chaco, HM: High Monte, SAY: Southern Andean Yungas, LM: Low Monte, HP: Humid Pampa; AP: annual precipitation, MAT: mean annual temperature.

**APPENDIX S2.**
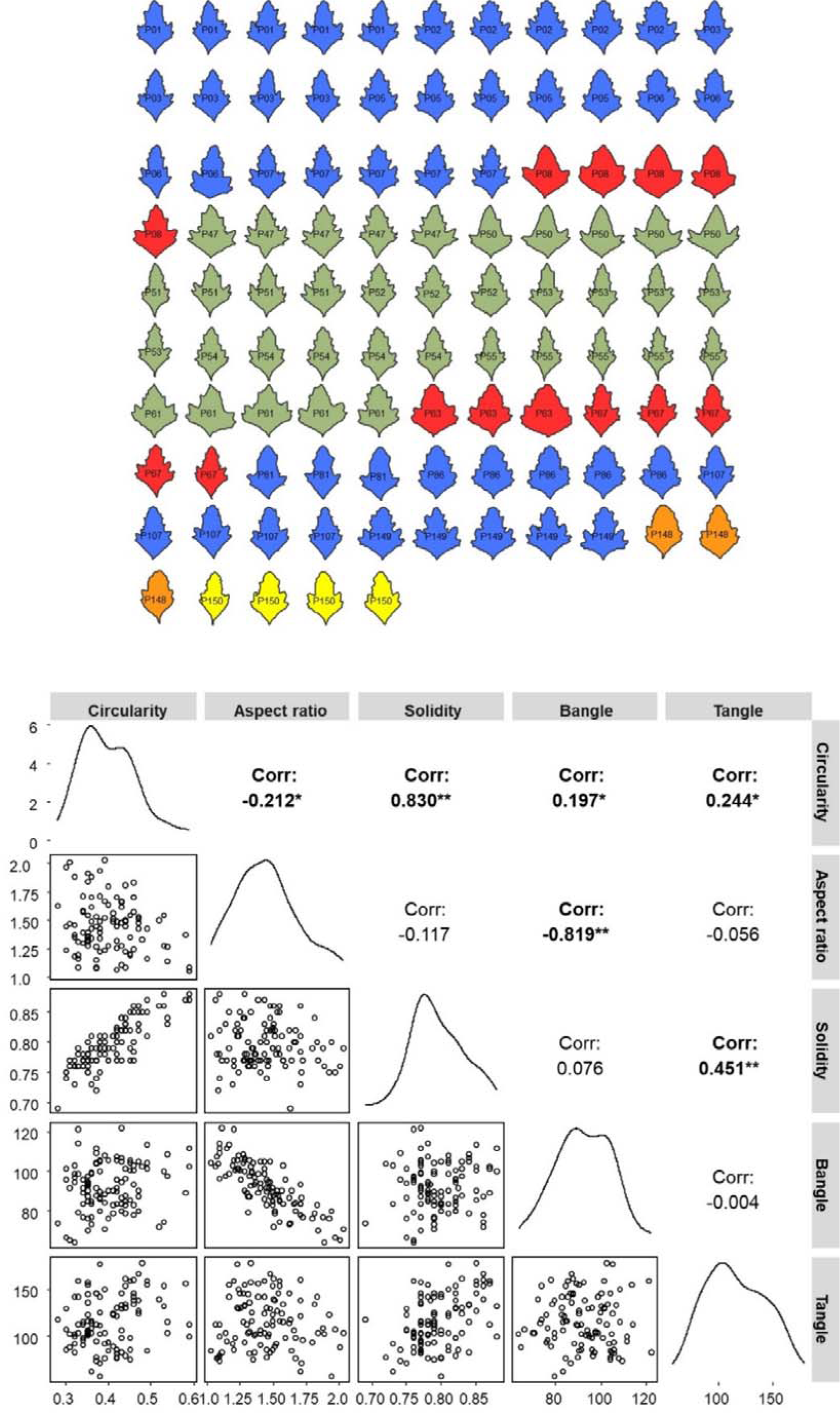
Leaf shape diversity in Argentinean avian goosefoot populations relationships among shape descriptors. Code inside each leaf (upper) is associated with population and color with ecoregions: blue (Dry Chaco), red (Southern Andean Yungas), green (High Monte), orange (Low Monte) and yellow (Humid Pampa). The scatterplot matrix (below) of the associations among shape descriptors is shown, bold letters indicate significant correlation.

**APPENDIX S3.**
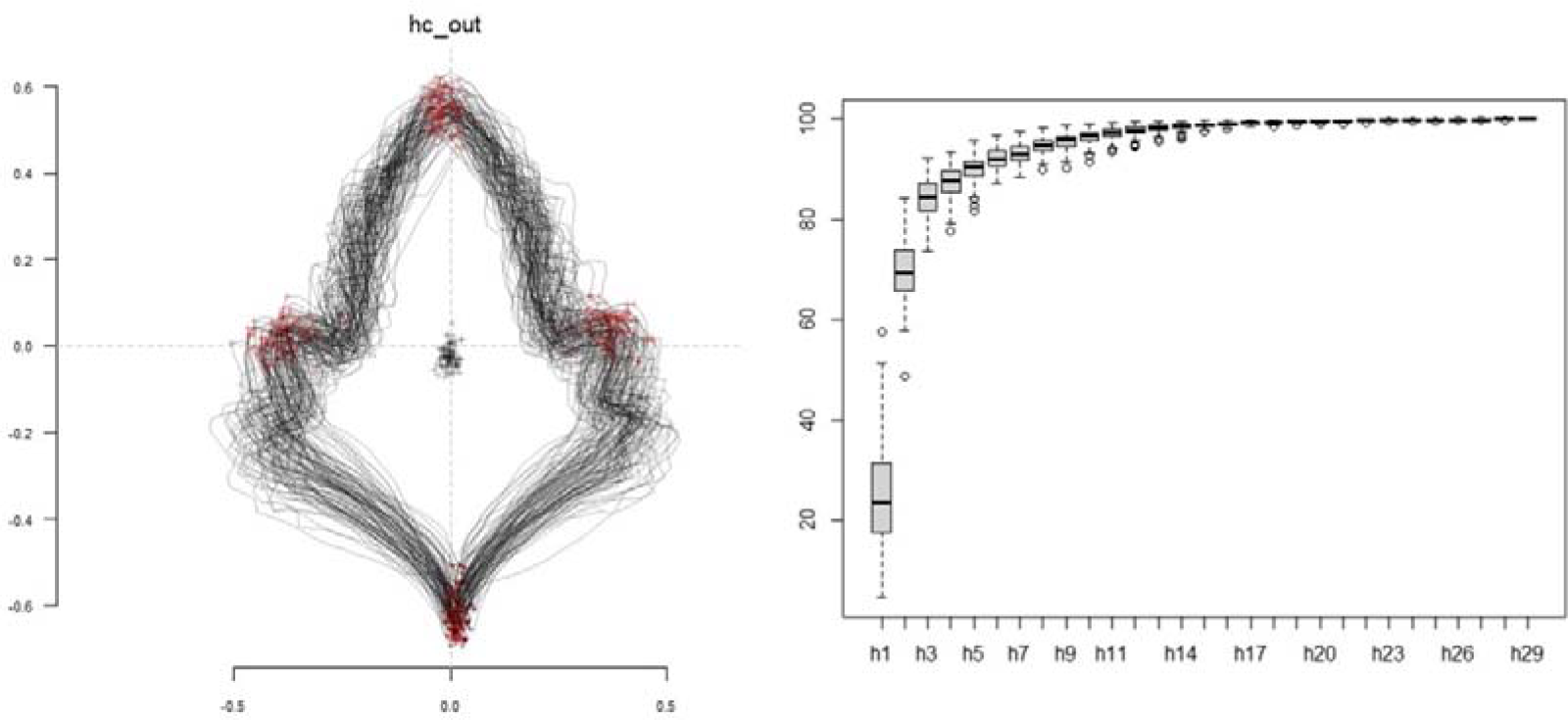
Aligned outlines of avian goosefoot leaves generated by Generalized Procrustes Analysis (left) and the number of harmonics retained for EFDs analysis (right).

**APPENDIX S4.**
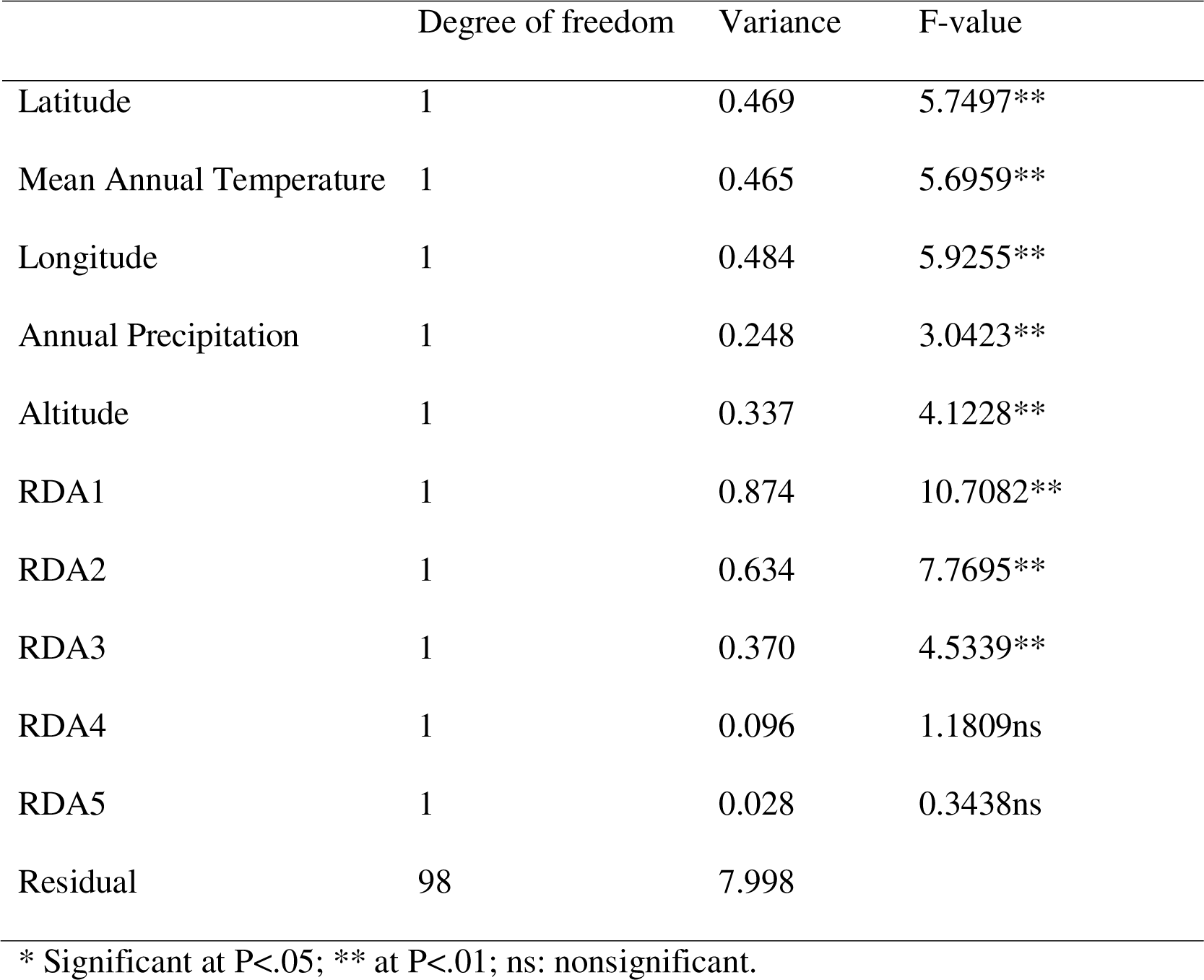
ANOVA results for predictors variables and the first five axes of Redundancy Analysis (RDA)

## REFERENCES

Bertero, H. D. 2021. Chapter 7 - Quinoa. In V. O. Sadras, and D. F. Calderini [eds.], Crop Physiology Case Histories for Major Crops, 250–281. Academic Press.

Bohra, A., B. Kilian, S. Sivasankar, M. Caccamo, C. Mba, S. R. McCouch, and R. K. Varshney. 2022. Reap the crop wild relatives for breeding future crops. Trends in Biotechnology 40: 412–431.

Bonhomme, V., S. Picq, C. Gaucherel, and J. Claude. 2014. Momocs: Outline Analysis Using R. Journal of Statistical Software 56: 1–24.

Curti, R. N., P. Ortega-Baes, S. Ratto, and D. Bertero. 2022. Harnessing phenological traits of wild ancestor *Chenopodium hircinum* to improve climate adaptation of quinoa. Crop and Pasture Science 73: 1058–1068.

Curti, R. N., P. Ortega-Baes, J. Sajama, D. Jarvis, E. Jellen, M. Tester, and D. Bertero. 2023. Exploration and Collection of Quinoa’s Wild Ancestor in Argentina. In R. Choukr-Allah, and R. Ragab [eds.], Biosaline Agriculture as a Climate Change Adaptation for Food Security, 167–178. Springer International Publishing, Cham.

Dryden, I. L., and K. V. Mardia. 2016. Statistical Shape Analysis: With Applications in R. John Wiley & Sons.

Fritz, M. A., S. Rosa, and A. Sicard. 2018. Mechanisms Underlying the Environmentally Induced Plasticity of Leaf Morphology. Frontiers in Genetics 9. 478. 10.3389/fgene.2018.00478

Frye, H. A., K. Mocko, T. E. Moore, C. D. Schlichting, and C. S. Jones. 2020. Leaf margins in a deciduous lineage from the Greater Cape Floristic Region track climate in unexpected directions. American Journal of Botany 107: 735–748.

Givnish, T. J., and R. Kriebel. 2017. Causes of ecological gradients in leaf margin entirety: Evaluating the roles of biomechanics, hydraulics, vein geometry, and bud packing. American Journal of Botany 104: 354–366.

Gleason, S. M., C. J. Blackman, S. T. Gleason, K. A. McCulloh, T. W. Ocheltree, and M. Westoby. 2018. Vessel scaling in evergreen angiosperm leaves conforms with Murray’s law and area-filling assumptions: implications for plant size, leaf size and cold tolerance. New Phytologist 218: 1360–1370.

Hinojosa, L., J. A. González, F. H. Barrios-Masias, F. Fuentes, and K. M. Murphy. 2018. Quinoa Abiotic Stress Responses: A Review. Plants 7: 106.

Hightower, A. T., D. H. Chitwood, and E. B. Josephs. 2024. Herbarium specimens reveal links between Capsella bursa-pastoris leaf shape and climate. bioRxiv: 2024.02.13.580180.

International, B., F. para la P. e I. de P. Andinos, I. N. de I. A. y Forestal, I. F. for A. Development, and F. and A. O. of the U. Nations. 2013. Descriptors for quinoa (Chenopodium quinoa Willd.) and wild relatives. Bioversity International.

Jarvis, D. E., Y. S. Ho, D. J. Lightfoot, S. M. Schmöckel, B. Li, T. J. A. Borm, H. Ohyanagi, et al. 2017. The genome of *Chenopodium quinoa*. Nature 542: 307–312.

Lesjak, J., and D. F. Calderini. 2017. Increased Night Temperature Negatively Affects Grain Yield, Biomass and Grain Number in Chilean Quinoa. Frontiers in Plant Science 8.

Li, Y., D. Zou, N. Shrestha, X. Xu, Q. Wang, W. Jia, and Z. Wang. 2020. Spatiotemporal variation in leaf size and shape in response to climate. Journal of Plant Ecology 13: 87–96.

Mandák, B., K. Krak, P. Vít, Z. Pavlíková, M. N. Lomonosova, F. Habibi, L. Wang, et al. 2016. How genome size variation is linked with evolution within *Chenopodium* sensu lato. Perspectives in Plant Ecology, Evolution and Systematics 23: 18–32.

Maughan, P. J., L. Chaney, D. J. Lightfoot, B. J. Cox, M. Tester, E. N. Jellen, and D. E. Jarvis. 2019. Mitochondrial and chloroplast genomes provide insights into the evolutionary origins of quinoa (*Chenopodium quinoa* Willd.). Scientific Reports 9: 185.

Maxted, N., S. Kell, B. Ford-Lloyd, E. Dulloo, and Á. Toledo. 2012. Toward the Systematic Conservation of Global Crop Wild Relative Diversity. Crop Science 52: 774–785.

Nicotra, A. B., A. Leigh, C. K. Boyce, C. S. Jones, K. J. Niklas, D. L. Royer, and H. Tsukaya. 2011. The evolution and functional significance of leaf shape in the angiosperms. Functional Plant Biology 38: 535–552.

Oksanen, J., G. L. Simpson, F. G. Blanchet, R. Kindt, P. Legendre, P. R. Minchin, R. B. O’Hara, et al. 2001. vegan: Community Ecology Package. 2.6–6.1.

Olson, D. M., E. Dinerstein, E. D. Wikramanayake, N. D. Burgess, G. V. N. Powell, E. C. Underwood, J. A. D’amico, et al. 2001. Terrestrial Ecoregions of the World: A New Map of Life on Earth: A new global map of terrestrial ecoregions provides an innovative tool for conserving biodiversity. BioScience 51: 933–938.

Oyarzabal, M., J. Clavijo, L. Oakley, F. Biganzoli, P. Tognetti, I. Barberis, H. M. Maturo, et al. 2018. Vegetation units of Argentina. Ecología Austral 28: 40–63.

Peppe, D. J., D. L. Royer, B. Cariglino, S. Y. Oliver, S. Newman, E. Leight, G. Enikolopov, et al. 2011. Sensitivity of leaf size and shape to climate: global patterns and paleoclimatic applications. New Phytologist 190: 724–739.

Presotto, A., M. Cantamutto, M. Poverene, and G. Seiler. 2009. Phenotypic diversity in wild *Helianthus annuus* from Argentina. Helia 32: 37–49.

Renzi, J. P., C. J. Coyne, J. Berger, E. von Wettberg, M. Nelson, S. Ureta, F. Hernández, et al. 2022. How Could the Use of Crop Wild Relatives in Breeding Increase the Adaptation of Crops to Marginal Environments? Frontiers in Plant Science 13: 886162.

Royer, D. L., D. J. Peppe, E. A. Wheeler, and Ü. Niinemets. 2012. Roles of climate and functional traits in controlling toothed vs. untoothed leaf margins. American Journal of Botany 99: 915–922.

Royer, D. L., and P. Wilf. 2006. Why Do Toothed Leaves Correlate with Cold Climates? Gas Exchange at Leaf Margins Provides New Insights into a Classic Paleotemperature Proxy. International Journal of Plant Sciences 167: 11–18.

Royer, D. L., P. Wilf, D. A. Janesko, E. A. Kowalski, and D. L. Dilcher. 2005. Correlations of climate and plant ecology to leaf size and shape: potential proxies for the fossil record. American Journal of Botany 92: 1141–1151.

Ruiz, R. A., and H. D. Bertero. 2008. Light interception and radiation use efficiency in temperate quinoa (*Chenopodium quinoa* Willd.) cultivars. European Journal of Agronomy 29: 144–152.

SAS (SAS Institute, Inc.). (2004). SAS/STAT® 9.4: User’s Guide

Traiser, C., S. Klotz, D. Uhl, and V. Mosbrugger. 2005. Environmental signals from leaves – a physiognomic analysis of European vegetation. New Phytologist 166: 465–484.

Walters, T. W. 1988. Relationship Between Isozymic and Morphologic Variation in the Diploids *Chenopodium fremontii*, *C. neomexicanum*, C. palmeri, and C. watsonii. American Journal of Botany 75: 97–105.

Wang, M., S. M. Kornblau, and K. R. Coombes. 2018. Decomposing the Apoptosis Pathway Into Biologically Interpretable Principal Components. Cancer Informatics 17: 1176935118771082.

Westerband, A. C., J. L. Funk, and K. E. Barton. 2021. Intraspecific trait variation in plants: a renewed focus on its role in ecological processes. Annals of Botany 127: 397–410.

Wilson, H. D. 1988. Allozyme Variation and Morphological Relationships of *Chenopodium hircinum* (s.l.). Systematic Botany 13: 215–228.

Wilson, H. D. 1981. Genetic Variation among South American Populations of Tetraploid *Chenopodium* sect. Chenopodium subsect. Cellulata. Systematic Botany 6: 380.

Wilson, H. D. 1990. Quinua and Relatives (*Chenopodium* sect. *Chenopodium* subsect. *Celluloid*). Economic Botany 44: 92.

Yang, J., R. A. Spicer, T. E. V. Spicer, N. C. Arens, F. M. B. Jacques, T. Su, E. M. Kennedy, et al. 2015. Leaf form–climate relationships on the global stage: an ensemble of characters. Global Ecology and Biogeography 24: 1113–1125.

Young, L. A., P. J. Maughan, D. E. Jarvis, S. P. Hunt, H. C. Warner, K. K. Durrant, T. Kohlert, et al. 2023. A chromosome[scale reference of *Chenopodium watsonii* helps elucidate relationships within the North American A[genome *Chenopodium* species and with quinoa. The Plant Genome 16: e20349.

